# Enabling community-based metrology for wood-degrading fungi

**DOI:** 10.1101/815852

**Authors:** Rolando Perez, Marina Luccioni, Nathaniel Gaut, Finn Stirling, Rohinton Kamakaka, Katarzyna P. Adamala, Pamela A. Silver, Drew Endy

## Abstract

Lignocellulosic biomass could support a greatly-expanded bioeconomy. Current strategies for using biomass typically rely on single-cell organisms and extensive ancillary equipment to produce precursors for downstream manufacturing processes. Alternative forms of bioproduction based on solid-state fermentation and wood-degrading fungi can enable more direct means of manufacture. However, such practices are often *ad hoc* and not readily reproducible. We sought to develop standard reference strains, substrates, measurements, and methods sufficient to begin to enable reliable reuse of mycological materials and products. Specifically, we show that a widely-available and globally-regularized consumer product (Pringles™) can support the growth of wood-degrading fungi, and that growth on Pringles™ can be correlated with growth on a fully-traceable and compositionally characterized substrate (NIST Reference Material 8492 Eastern Cottonwood Biomass). So established, five laboratories were able to compare measurements of wood-fungus performance via a simple radial extension growth rate assay. Reliable reuse of materials, measures, and methods is necessary to enable distributed bioproduction processes that can be adopted at all scales, from local to industrial.

## INTRODUCTION

The contiguous United States (U.S.) contains approximately 700 million tons of potential lignocellulosic biomass^1^. The available biomass is generally a mixture of forest and agricultural residues, urban organic waste, and algae. Bioproducts derived from lignocellulosic biomass, such as chemicals, enzymes, bioplastic bottles and packaging, and textiles contributed an estimated $393 billion and 4.2 million jobs to the U.S. economy in 2014^2–4^. The U.S. Department of Agriculture (USDA) reports that global revenues from bio-based chemicals alone will exceed $500 billion dollars per year by 2025^5^. Further development of infrastructure for utilizing distributed biomass resources could help enable a “Bio-Belt” that revitalizes rural manufacturing and benefits historically-disenfranchised communities^6^.

Current biomanufacturing processes typically focus on microbial conversion of lignocellulosic biomass feedstocks to sugars by way of enzymatic hydrolysis followed by fermentation into higher-value products^7^. Such approaches require resource-intensive biomass preprocessing, liquid bioreactors, and ancillary equipment^2^. In contrast, wood-degrading filamentous fungi – evolved to grow directly from lignocellulose – could provide a more straightforward means of manufacture that is less resource intensive and expands what bioproducts can be made^8–11^. The direct use of wood-degrading filamentous fungi could also facilitate a more ‘circular’ economy supporting sustainable food production, manufacturing, and energy generation^12^. Direct use of wood-degrading fungi could even enable new opportunities for innovation in the global economy, especially in developing countries^12–14^.

Fungi form diverse relationships with viruses, algae, plants, animals, insects, and bacteria, and are critical to human health^15–21^. Filamentous fungi, in particular, are distinguished by their multinucleate cells, known as hyphae, chitinous-composite cell walls, vast repertoire of degradative enzymes and secondary metabolites, and interconnected networks of cells, known as mycelium, that form through a combination of apical growth, hyphal branching, and regulated self-fusion^22–31^. Growing mycelium penetrate and degrade the substrate upon which the organism lives and uptake and shuttle nutrients throughout complex hyphal networks^23^.

Mushrooms, the reproductive structures of wood-degrading filamentous fungi, are widely-used as foodstuffs and as a source of medicines and are consequentially revered by many cultures^30–32^. Additionally, mycological materials made from the tissues of mushrooms are used in meat replacements, filtration and remediation of contaminated water, packaging materials, furniture, textiles for the fashion industry, architectural design, art, materials for super capacitors and batteries, anti-viral therapeutics for bees, and nanoparticle synthesis^35, 36, 45, 46, 37–44^. Most commercial mycological materials are fabricated by placing an organism and substrate mixture into a preformed mold and allowing the mycelium to process the substrate via solid-state fermentation. The process itself is typically halted via thermal inactivation followed by downstream processing.

Recent studies have elucidated the impacts that choice of organism, genetic modification, substrate composition, and environmental conditions have on the macroscopic properties of mycelium-based materials^47–54^. Most organisms now used to produce mycelium materials are of the class Agaricomycete in the division Basidiomycota. The Agaricomycete are less-extensively studied and developed as some of their Ascomycota counterparts, such as *Saccharomyces cerevisiae*, *Neurospora crassa*, or A*spergillus niger*^55–57^. The model mushrooms that have been advanced in academic research, such as *Coprinopsis cinera* for mushroom genetics, *Agaricus bisporus* for food, *Schizophyllum commune* for mushroom morphogenesis, and *Phanerochaete chrysosporium* for lignocellulosic biomass degradation, are different from the organisms that are used by industry, such as *Ganoderma lucidum* and *Trametes versicolor*^11, 58–66^. Additionally, the substrates that industry groups use in production are often proprietary recipes that are seldom shared. However, substrate composition has been demonstrated to be critical in defining the macroscopic properties of mycological materials. For example, Elsacker et al. demonstrated the use of standard methods for preparing various lignocellulosic substrates only to find that substrate physical characteristics (e.g., fiber type) may have a greater impact on macroscopic material properties than the chemical composition of the substrate itself^67^. Elsacker et al.’s work illustrates the need for standards that facilitate the reliable reuse of materials and measures comprising mycelium-based fabrication processes.

Reliable reuse of materials and measures has long been recognized as essential for commerce and civilization^68–70^. For example, Section 8 of the U.S. Constitution grants Congress the power to, “fix the Standard of Weights and Measures”^71^. Standards are so useful that the U.S. Commerce Department operates the National Institute of Standards and Technology (NIST) to establish essential measures for the U.S. economy. Additionally, multiple industries support sector-specific standards-setting organizations including the International Electrotechnical Commission (IEC) for electrical and electronic engineering standards, the Internet Engineering Task Force (IETF) for information networking standards, and the International Organization for Standardization (ISO) for materials, products, processes, and services standards. Beginning in 1999, reuse of standard measures, materials, and methods began to prove useful for synthetic biology; but, the resulting impacts of standardization with and of living matter have not yet matched other technology-enabled sectors^72–79^. Although humans have long-partnered with filamentous fungi there are yet to emerge any standards supporting the use and reuse of wood-degrading fungi in mycologically-based bioproduction^80–83^.

Here, we sought to develop materials, measures, and methods that begin to support community-based metrology for wood-degrading fungi. Specifically, we sourced sequenced strains of wood-degrading fungi and used standard plate-based colony radial extension assays to determine strain performance^84, 85^. We show that growth performance of fungi on Pringles™ correlates well with performance on a fully characterized and traceable NIST reference material. We also demonstrate via interlaboratory experimentation that measurements of wood-fungi performance can be coordinated across many locations.

## RESULTS

Most mycelium-based fabrication processes use organisms whose genomes have not been sequenced. Going forward, working with sequenced organisms is absolutely critical to support science- and technology-enabled improvements in strains and manufacturing processes. Thus, in getting started, we worked to source five sequenced strains of industrially-relevant wood-degrading fungi: *Pleurotus ostreatus,* an important gourmet mushroom that is used in mycelium-materials; *S. commune,* a widely-recognized model organism for mushroom morphogenesis; *T. versicolor,* a medicinal mushroom used for remediation and mycelium materials; *G. lucidum*, a medicinal mushroom and favorite of the nascent mycelium materials industry, and *P. chrysosporium*, a recognized model organism for processing lignocellulosic biomass.

Obtaining sequenced strains for each selected organisms was complicated. For example, we first acquired a sample of *P. ostreatus* (strain ID PC-9) from the Spanish Type Culture Collection (CECT). Acquiring samples of *P. ostreatus* from CECT required completing an ad hoc materials transfer agreement, USDA APHIS permitting, Spanish Customs export compliance requirements which included affirming future compliance with the Convention on Biological Diversity and the Nagoya Protocol, and finally U.S. Customs import requirements. However, upon first use of the so-sourced sample, we observed microbial contamination in our stock cultures, which we were able to clear through subsequent sub-culture. We experienced similar challenges acquiring sequenced strains of *G. lucidum* from various sources. We eventually acquired *G. lucidum* (strain ID: 10597-SSI)*, T. versicolor* (strain ID: FP101664-Spp)*, P. chrysopsporium* (strain ID: RP-78) from the Reference Culture Collection at the Center for Forest Mycology at the USDA Forest Service Forest Products Lab. All strains had been sequenced by the U.S. Department of Energy Joint Genome Institute (JGI), and the genome assemblies of each strain can be accessed via the JGI MycoCosm portal (mycocosm.jgi.doe.gov) (Table S1). Importantly, all organisms as acquired are not on the U.S. Regulated Plant Pest list, which makes reuse simpler.

We then worked to confirm a suitable measurement method that can be readily used by a diversity of practitioners. First, we compared total biomass accumulation in liquid culture with plate-based measurements of colony radial extension to determine if either method would be practical for community-based measurements. Liquid-culture methods do not resemble the ambient conditions encountered in solid-state fermentation, typically take more than five days to complete, use sample volumes greater than 10 mL, and require ancillary equipment to properly culture the samples, harvest and dry the biomass, and measure the dry cell weight. In our small pilot studies, we found 5 mL culture volumes of various fungi and substrate combinations resulted in irreproducible results, and so we abandoned a liquid culture-based measurement method in supporting distributed mycometrology.

In contrast, plate-based colony radial extension measurement methods more closely reflect the ambient conditions of solid-state fermentation processes and require less equipment than liquid-based methods. Our use of the plate-based colony radial extension rate measurement method consists of daily traces of the colony radius, followed by imaging of the traces, and image analysis to quantify extension rates (Figure 1). In addition to quantifying colony radial extension rates, we observed qualitative differences in hypha density and hyphal branch rate between various fungi and substrate combinations (Figure S1). The plate-based assay is sensitive to error in daily measurements, especially with slow growing colonies on unfavorable substrates, and also to microbial contamination during prolonged incubation periods on substrates that lack antibiotics. Nevertheless, plate-based colony radial extension rate measurements provide an accessible metrological method that can be used by many practitioners.

**Figure 1.**
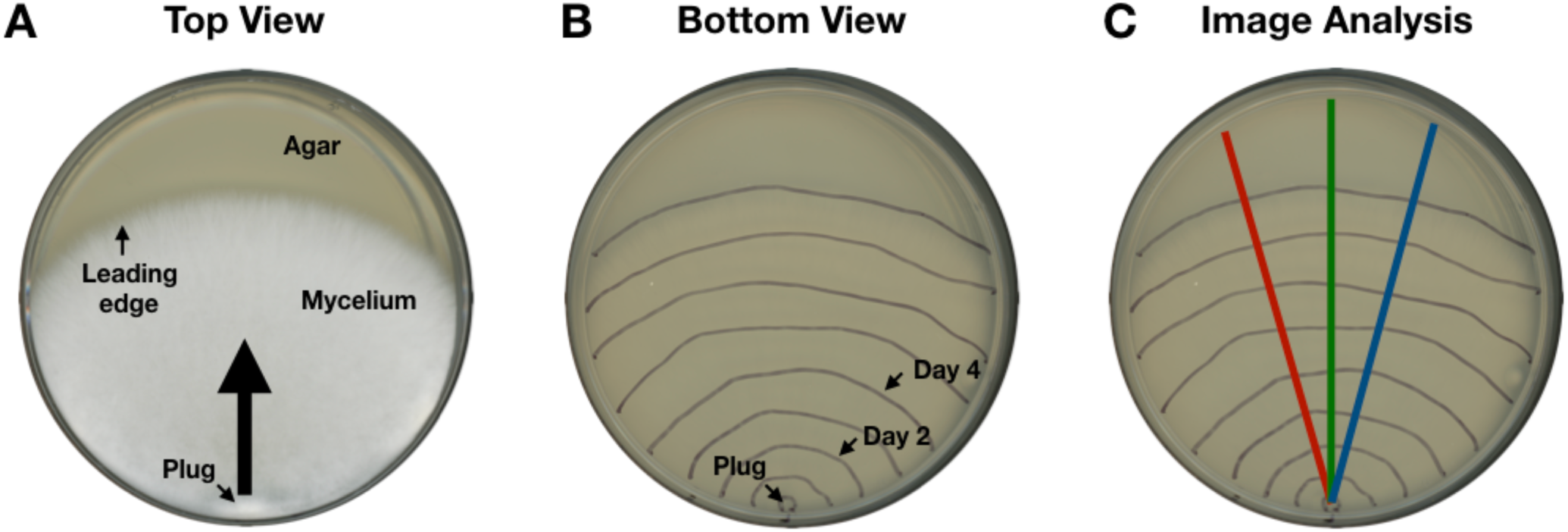
Plate-based radial extension measurements are an accessible method for quantifying the performance of wood-degrading fungi. (A) The fungal colony is started from a single inoculation point of fungal tissue. The 5 mm tissue plug is taken from a common source plate of fungal tissue, applied to an experimental plate, and incubated in darkness at 30C; resulting in leading edge radial growth along the agar surface in the direction indicated by the large black arrow. (B) The leading-edge of the growing colony is traced every 24 hours with a marker. (C) On the final day traces are imaged and analyzed using ImageJ. Measurements are made along each axis, red, green, and blue at intersections with the leading edge traces.

Wood-fungus are often grown on “found” feedstocks that can be sourced locally. To better understand the range of growth performance as a function of variation in feedstocks we measured fungal colony growth rates for *G. lucidum, T. versicolor,* and *P. chrysosporium* on agar-based substrates made from aqueous extracts of five locally-available substrates: plant compost from the Stanford Farm, horse manure from the Stanford Barn, a commercially-available wood chip, laboratory-grade potato dextrose yeast extract (PDYA), and a yeast synthetic defined media (YSD) (Table S2). We found that all fungi tested grew on all substrates (Figure 2). However, extension rates varied up to 3-fold across organisms and up to 7.5-fold across substrates. *P. chrysosporium* had the greatest average extension rate followed by *T. versicolor,* and *G.* lucidum.

**Figure 2.**
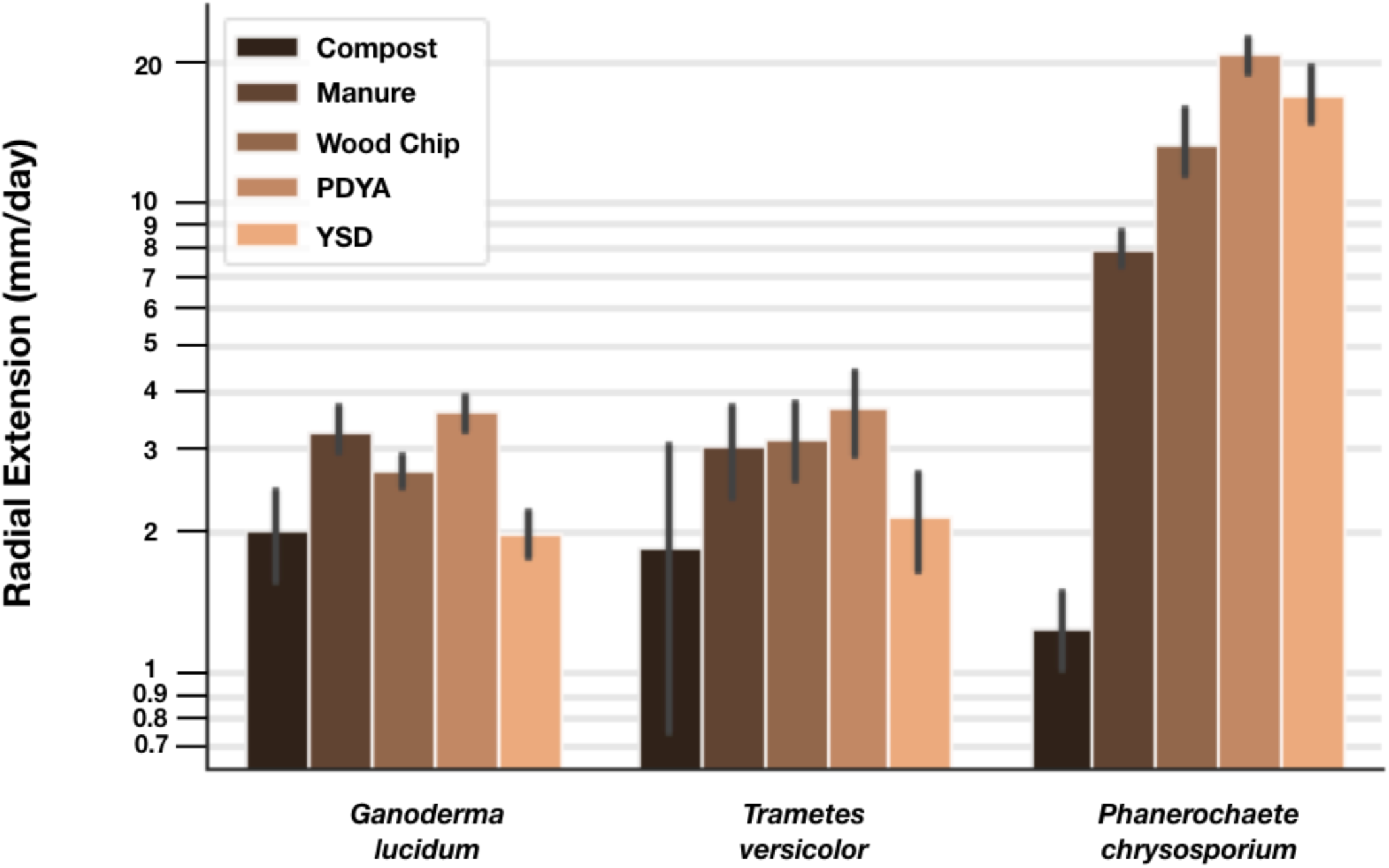
Extension rates of selected wood-degrading fungi vary up to 7.5-fold with respect to substrate composition. We measured radial extension rates of three different sequenced wood-degrading fungi on six different substrates. *P. chrysosporium* had the greatest average extension rate followed by *T. versicolor,* and *G.* lucidum. Rates below 2 mm/day are shown but are considered to be at the limit of our measurement technique. Statistically significant differences were detected by way of Tukey’s HSD (p<0.05) for *P. chrysosporium* and *T. versicolor*, and *P. chrysosporium* and *G. lucidum.* Statistically significant differences were detected by way of Tukey’s HSD (p<0.05) for Compost and Wood Chip, Manure PDYA, and YSD; between Manure and PDYA; and between PDYA and Wood Chip. The reported rates and error bars are the mean and standard deviation of biological replicates, respectively; n=6.

Differences in growth among strains and across feedstocks make it difficult to compare measures of performance across locations and over time. One impractical approach to enabling comparable measurements would be to regularize all feedstocks and strains. Instead we sought to identify a reference material that could be used as a standard substrate for calibrating wood fungus growth. Practitioners might make measurements of growth on both the standard and local substrates, allowing for comparison of performance in relation to a standard substrate used in common. For example, Eastern Cottonwood Whole Biomass Feedstock (NIST RM 8492) was developed as a standard lignocellulosic reference material supporting biomass research. RM 8492 has undergone full compositional characterization, has been used in NIST-led interlaboratory studies, and is available for purchase as a calibration tool for biomass feedstock production facilities (Table S3). However, the material is only available in limited quantities and expensive ($8.26/gram).

We thus sought standard substrates that are low-cost, can be sourced by anyone anywhere, and, if possible, related to NIST RM 8492. To do so we sourced materials that might support wood-fungus growth and that can be found throughout global consumer-product supply chains. Specifically, we sourced cardboard from Amazon Prime™ shipping boxes, wood chips from hard- and soft-wood shipping pallets, and Pringles™ – a consumer food product – as candidate standard substrate materials. To elaborate on the motivations guiding our sourcing process via one example: Pringles™ are a standardized potato-based food that is available in approximately 140 countries^86^; Pringles™ generated a reported $342 million in retail sales in the UK alone in 2018, a scale at which economic incentives ensure that Pringles™ remain available regardless of UK-EU politics^87^; Pringles™ themselves are composed primarily of potato starch and are produced to specification by only five factories globally (Malaysia, Belgium, China, Poland, U.S.); Pringles™ shipments typically arrive on standardized pallets and within standardized cardboard tubes.

We tested the performance of four industrially-relevant wood-degrading fungi – *S. commune*, *T. versicolor*, *G. lucidum*, and *P. chrysosporium –* on the candidate widely-available substrates. We observed that Pringles™ supported the greatest extension rate across all organisms tested (Figure 3). We also determined that extension rates varied up to 2.5-fold across the different organism and substrate combinations. *P. chrysosporium* had the greatest average extension rate, followed by *G. lucidum*, *T. versicolor*, and *S. commune.* For the four fungi tested, aqueous extracts of woodchips made from softwood pallets exhibited the slowest extension rates; rates below 2 mm per day are shown but are considered at the limit of our measurement technique.

**Figure 3.**
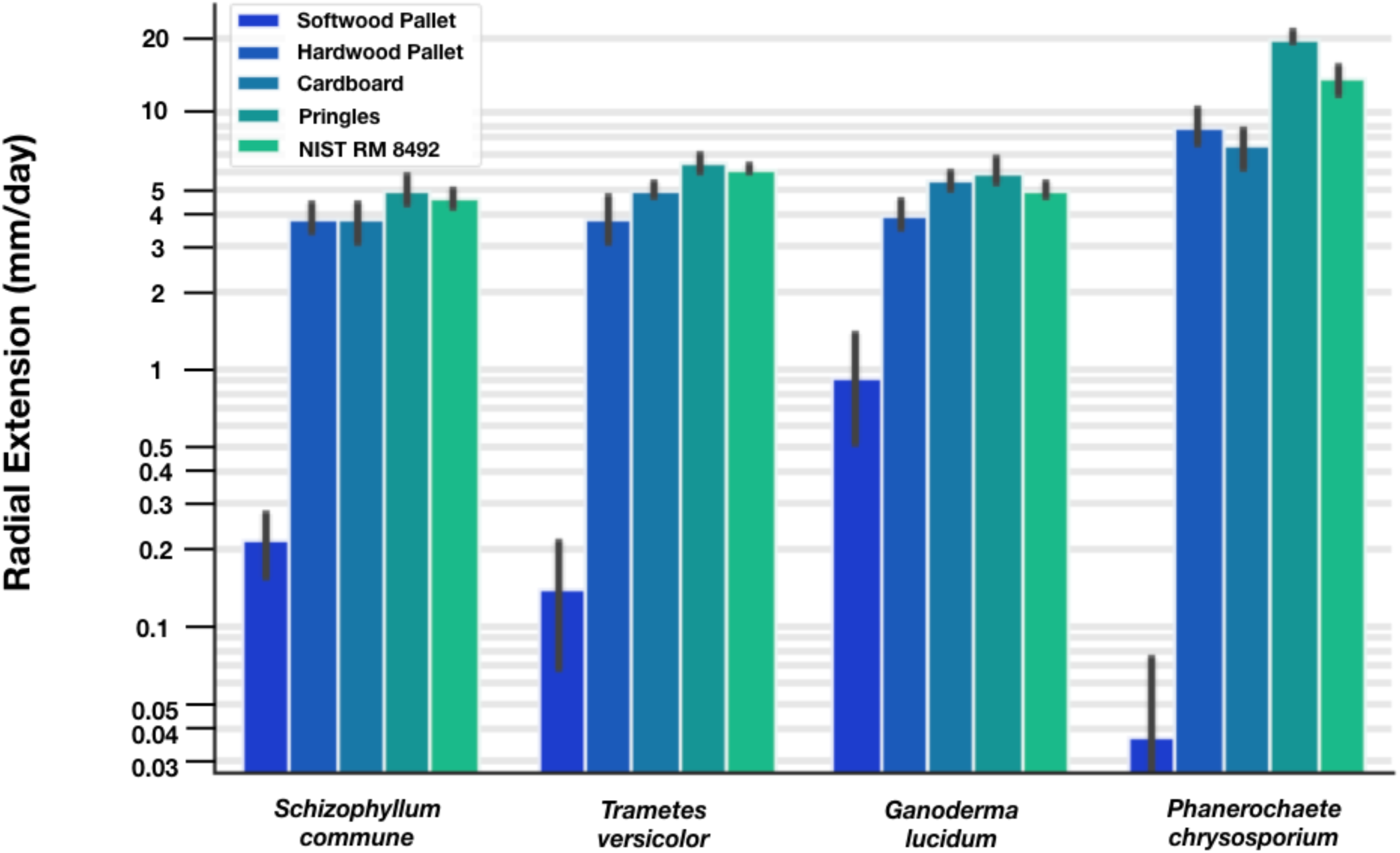
Pringles™ are a low-cost readily-available material supporting growth measurements of wood-degrading fungi. Extension rates vary up to 4-fold across various consumer materials. We measured radial extension rates of four select sequenced strains of industrially relevant wood-degrading fungi on four materials that can be found at the edges of global shipping networks. *P. chrysosporium* had the greatest average extension rate, followed by *G. lucidum*, *T. versicolor*, and *S. commune.* Rates below 2 mm/day are shown but are considered to be at the limit of our measurement technique. Statistically significant differences were detected by way of Tukey’s HSD (p<0.05) for *P. chrysosporium* and all other strains. Statistically significant differences were detected by way of Tukey’s HSD (p<0.05) for Cardboard and Softwood Pallet, Pringles™, and NIST RM 8492; between Hardwood Pallet and Softwood Pallet, Pringles™, and NIST RM 8492; between Pringles™ and NIST RM 8492, and Softwood Pallet; and NIST RM 8492 and Softwood Pallet. The reported rates and error bars are the mean and standard deviation, respectively (n=9).

We chose to proceed with Pringles™ as a widely-available substrate reference material primarily because it supported the most robust growth among all of the fungi tested, implying that Pringles™-based media may support the growth of a wider range of fungi. Though robust growth is not necessarily a prerequisite for substrate reference materials, the ability to support the growth of a diversity of fungi other than wood-degrading fungi could be more generally useful; Pringles™ are composed primarily of sugar-rich substances: dried potatoes, cornstarch, rice flour, and wheat starch, all of which are commonly-used substrates for mycological production. We then compared the performance of all organisms on media made from Pringles™ to performance on media made from NIST RM 8492 using a simple linear model (Figure 4). The observed correlation (R^2^ = 0.85) supports the potential to relate measurements of wood-fungus growth made by practitioners at the edges of consumer supply-chains to measurements made by well-resourced practitioners with access to fully-traceable and compositionally-characterized materials.

**Figure 4.**
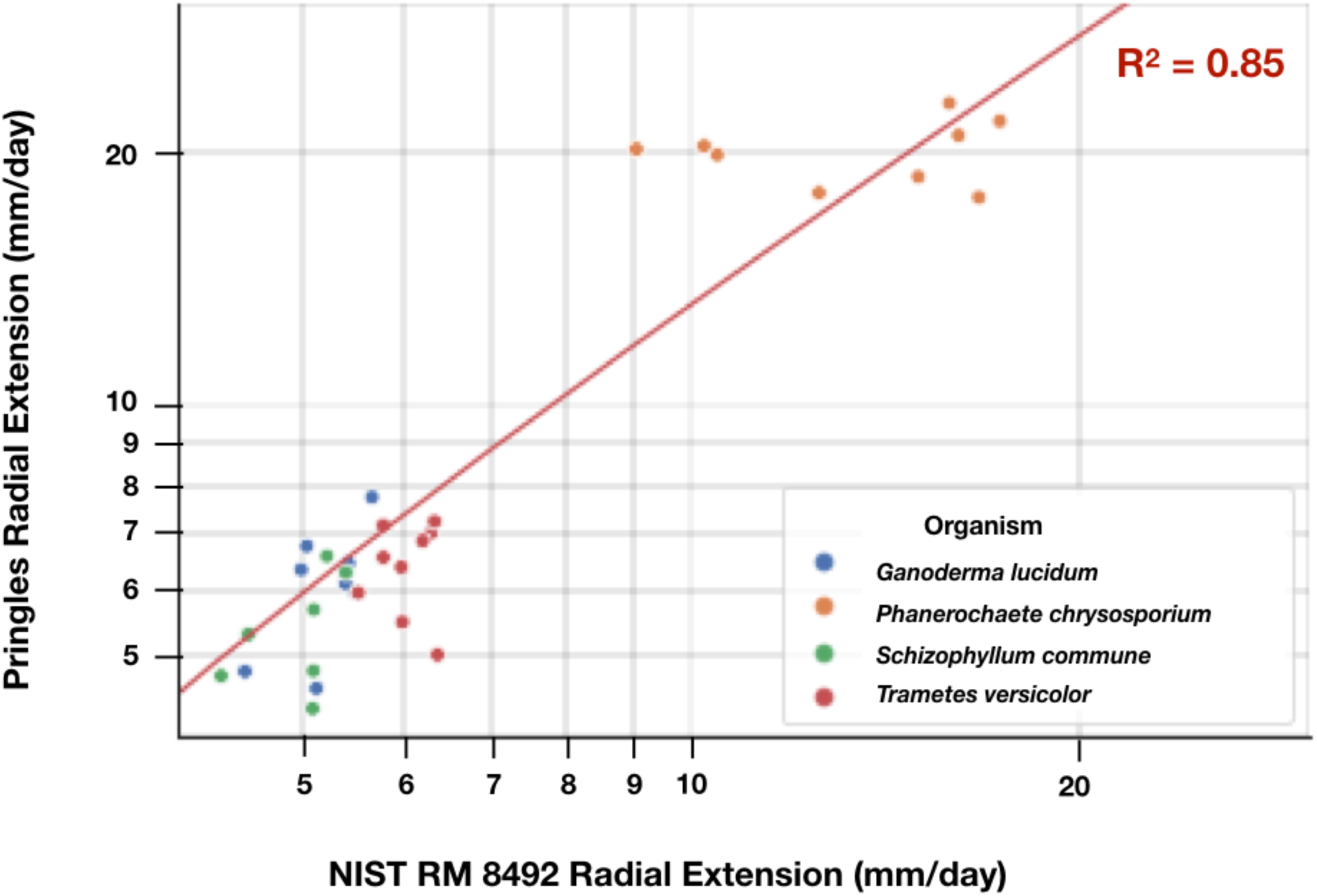
Pringles™ and NIST RM 8492 extension rates are well correlated (R^2^ = 0.85, n=9). We propose readily available consumer materials such as Pringles™ and compositionally characterized NIST RM 8492 as reference materials. Measurements made on NIST RM 8492 can be used to calibrate measurements of extension rates on compositionally characterized substrates with that of substrates made with standardized readily-available consumer materials. Such calibrations can support coordination between practitioners in a wide variety of resource settings.

To further explore variation in wood-fungus growth measurements as a function of Pringles™ sourcing and composition, we measured and compared the extension rates of *G. lucidum* on substrates made from Original-flavor Pringles™ sourced directly from China, Malaysia, Poland, Belgium, and the U.S., and also from substrates made from BBQ-, Sour Cream-, Cheddar-, Honey Mustard-, and Pizza-flavored Pringles™ sourced from the U.S. (Figure 5). Radial extension rates of *G. lucidum* on substrates sourced from the five Pringle™ production factories (Original flavor) and other Pringle™ flavors (assorted) varied ∼1.2-fold. Original-flavor Pringles™ made in China supported the highest extension rate while Honey Mustard-flavor supported the highest extension rate among the U.S.-made Pringle™ flavors tested; observed coefficients of variation were 0.12 and 0.04 for factories and flavors, respectively.

**Figure 5.**
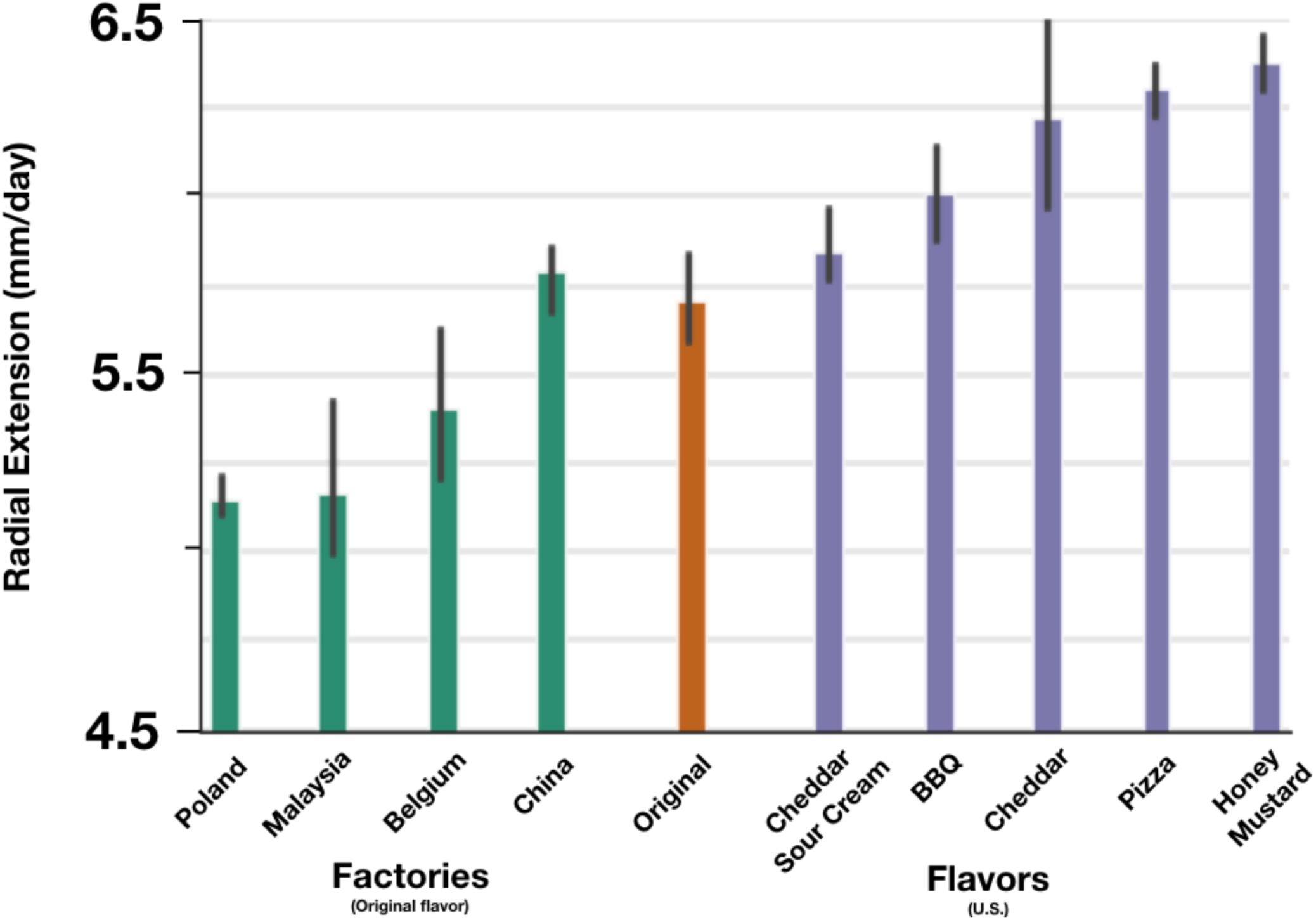
Extension rates on Original-flavor Pringles™ sourced from all five global factories and on assorted flavors from the U.S. vary less than 1.3-fold. We measured the extension rate of *Ganoderma lucidum* on substrates made from Original-flavor Pringles™ sourced from each of the five Pringles™ production factories as well as substrates made from BBQ-, Sour Cream-, Cheddar-, Honey Mustard-, and Pizza-flavored Pringles™. Original-flavor Pringles™ made in China supports the highest extension rate while Honey Mustard flavor supports the highest extension rate among the US-made Pringles™ flavors tested. No statistically significant differences were detected between production factories, flavors, or between factories and flavors (Tukey’s HSD, p<0.05) supporting their utility as standard reference materials. The reported rates and error bars are the mean and standard deviation, respectively. n=3 for all rates except for Original-USA, n=4. Replicates for two groups, Original (made in TN, USA) and USA (Original flavor, made in TN, USA), were combined into Original-USA and two plates per substrate were dropped due to microbial contamination.

**Figure 6.**
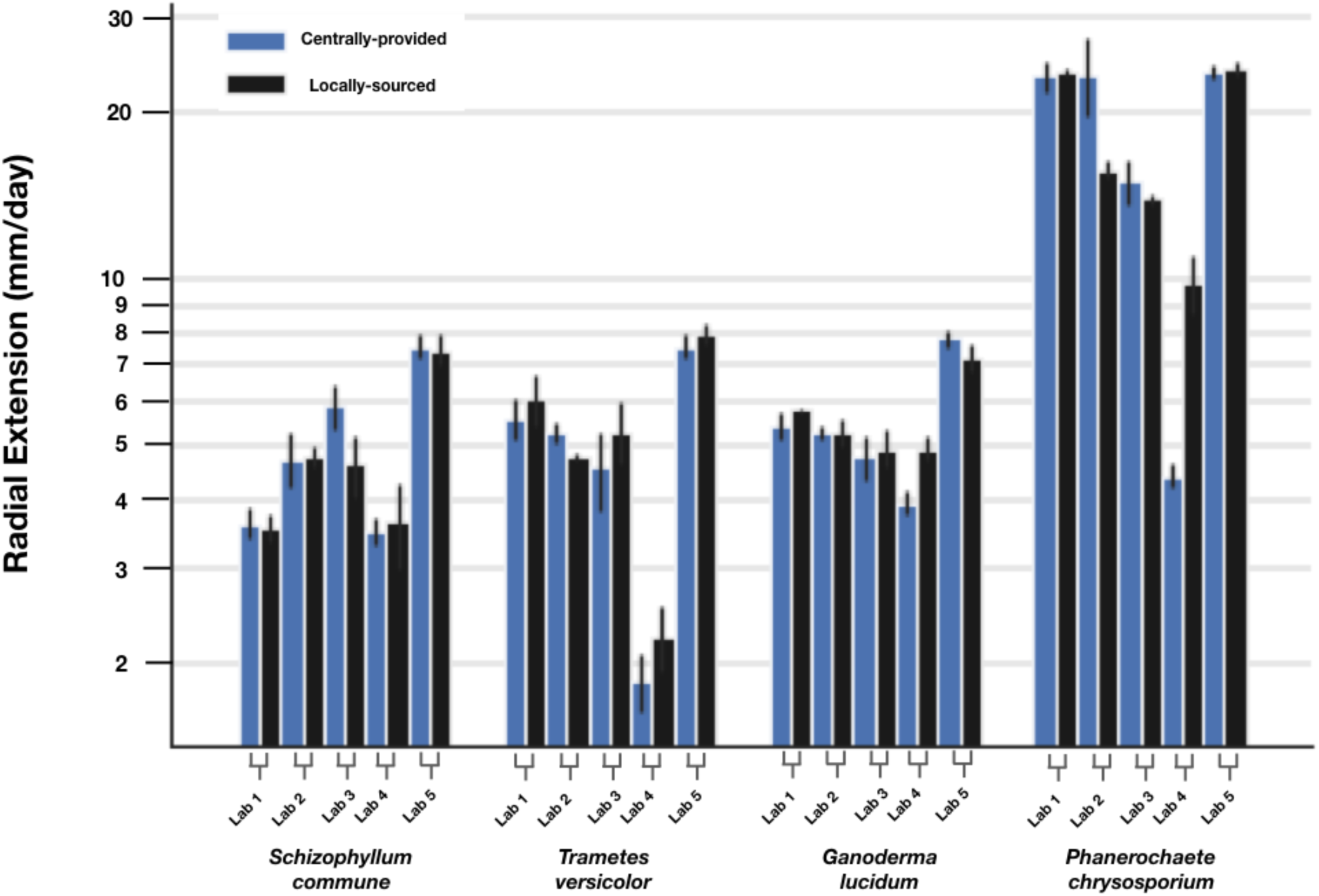
Community-based measurements of radial extension on media made from locally-sourced and centrally-provided Pringles™ are well matched. We distributed a measurement kit containing our proposed reference materials: *G. lucidum* (10597-SSI), *T. versicolor* (FP101664-Spp)*, S. commune* (4.8B)*, P. chrysosporium* (RP-78), and Original-flavor Pringles™. Participating labs also acquired their own local form of Original-flavor Pringles™ and used both substrate sources for radial extension experiments. Statistically significant differences per Tukey’s HSD (p<0.05) were detected for Lab 4 measurements of *P. chrysosporium* extension rates across locally-sourced and centrally-provided substrates. Statistically significant differences per Tukey’s HSD (p<0.05) were detected across labs when using centrally-provided Pringles™ between Lab 4 and Lab 5, Lab 4 and Lab 2, Lab 5 and Lab 3, and Lab 5 and Lab 2. Statistically significant differences per Tukey’s HSD (p<0.05) were detected across labs when using locally-sourced Pringles™ between Lab 4 and all labs (Lab 5, Lab 1, Lab 3, Lab 2), Lab 5 and Lab 3, and Lab 3 and Lab 2. The reported rates and error bars are the mean and standard deviation, respectively (n=3).

To practically explore the potential for reuse of consumer-sourced materials via a common measurement method we sent metrology kits to four participating laboratories (Figure S2). The kit included the four fungi strains of interest, a coring tool for excising fungal tissue plugs, cheese cloth for filtering aqueous extracts, centrally-provided Pringles™, and simple instructions for conducting colony radial extension measurements. Interlaboratory study participants independently acquired locally-sourced Pringles™, conducted experiments, and returned images of experimental plates for image processing and analysis. Measurements of growth rates on media made from locally-sourced versus centrally-provided Pringles™ substrates were within 1.5-fold, with the exception of Lab 4 measurements of *P. chrysosporium* growth rates. Observed coefficients of variation for growth rates obtained in our lab were 0.11, 0.04. 0.02, 0.16 for *S. commune*, *T. versicolor*, *G. lucidum*, and *P. chrysosporium*, respectively; coefficients of variation in growth rates across all other labs were 0.23, 0.28, 0.06, and 0.39 for *S. commune*, *T. versicolor*, *G. lucidum*, and *P. chrysosporium*, respectively

To explore reducing variation in measurements across participating laboratories we also calculated relative extension units (REU) for colony growth rates. In doing so we sought to leverage the use of relative units to normalize the performance of biological systems in relation to extrinsic factors that are difficult to identify or impossible to control, as has been previously established for reporting levels of gene expression^78^. Specifically, using growth rate data obtained with locally-sourced Pringles™ (Figure 5), we divided each lab’s reported extension rates for a particular organism by the lab’s reported extension rate for *G. lucidum* on locally-sourced Pringles™ (Figure 7). The resulting Relative Extension Units (REU) for each organism are within 2-fold with the exception of REU reported by Lab 4 for *T. versicolor* and *P. chrysosporium*. Reporting via REUs reduced coefficients of variation in all cases: by 80% for *G. lucidum*, by 33% for *S. commune*, by 34% for *T. versicolor*, and by 12% for *P. chrysosporium*.

**Figure 7.**
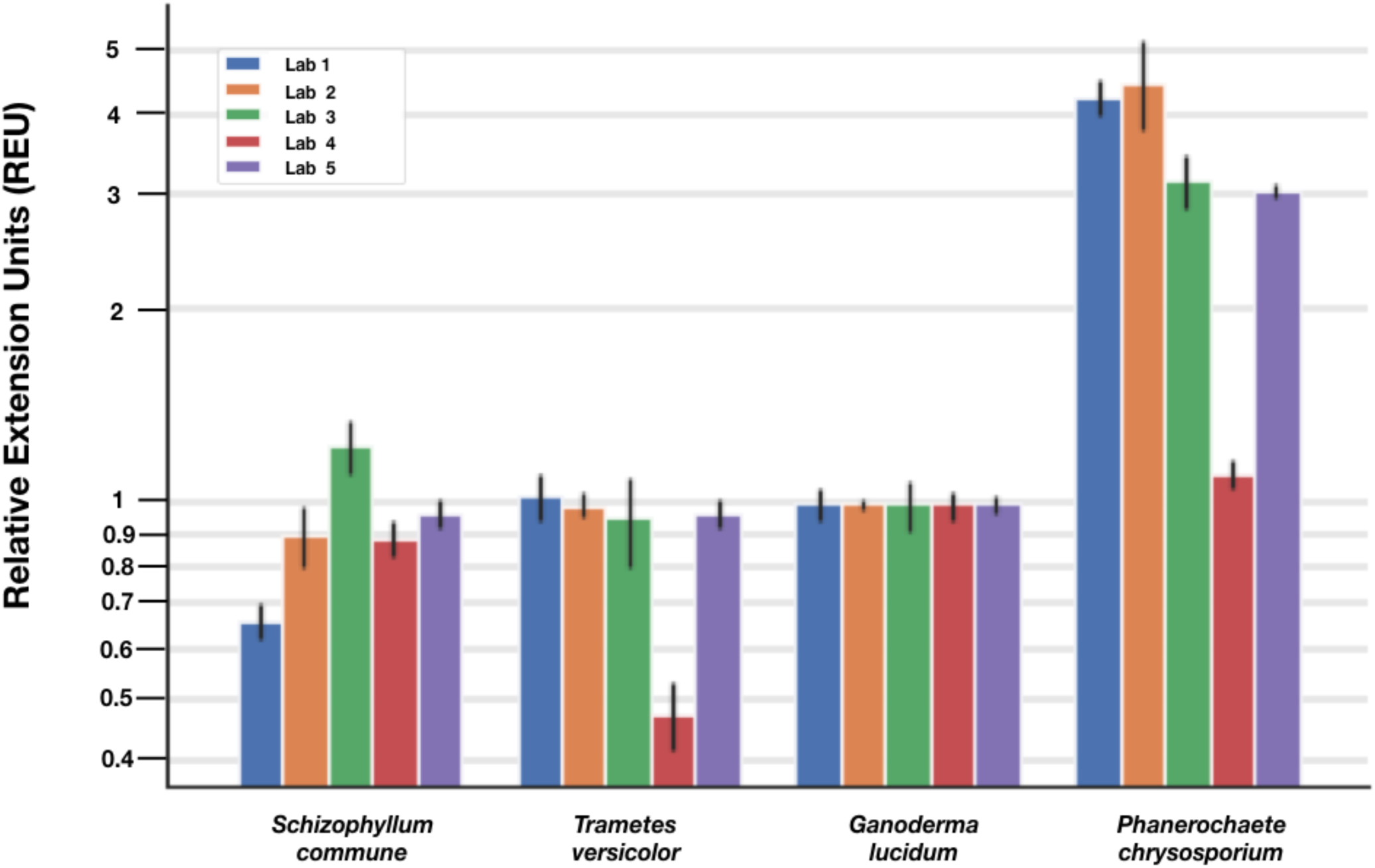
Reference materials and relative reference units improve community-based measurements of performance for wood-degrading fungi. Reported Relative Extension Units (REU) for each organism are within 2-fold with the exception of REU reported by Lab 4 for *T. versicolor* and *P. chrysosporium*. Using the same data from Figure 5, we divided each lab’s reported extension rates for a particular organism by the lab’s reported extension rate for *G. lucidum*. Reporting REUs reduced coefficients of variation for *G. lucidum* by 80%, for *S. commune* by 33%, for *T. versicolor* by 34%, and for *P. chrysosporium* by 12%. The reported rates and error bars are the mean and standard deviation of the normalized values, respectively (n=3).

## DISCUSSION

We established preliminary methods and materials that enable community-based metrology for wood-degrading fungi. We did so by sourcing sequenced organisms, identifying widely-available substrate feedstocks, and correlating growth performance to traceable reference materials via a simple measurement method. Taken together, these methods and materials should begin to improve the reproducibility and reliability of fabrication processes involving mycelium materials.

Specifically, we demonstrated that widely-available consumer materials, such as Pringles™, can serve as standard reference materials for mycological growth processes. As noted, Pringles™ can be found in most countries and are composed primarily of sugar-rich materials that are commonly used for mycological production. Furthermore, Pringles™ are ∼800-fold more affordable than fully characterized and traceable reference materials (NIST RM 8492 is $8.26/gram while Pringles™ are $0.01/gram). To be clear, Pringles™ themselves would likely not be a favored source of substrate for large-scale manufacturing processes; rather, Pringles™ can serve as an affordable and widely-available reference material that can be accessed and used by many practitioners across many locations.

We observed that flavored Pringles™ generally supported slightly faster growth than Original-flavor Pringles™ (Figure 5). Although we were not able to conduct compositional analysis of the flavored Pringles™ substrates, each product comes with a nutritional label and ingredients list. Many of the ingredients are the same across Pringles™ flavors, with Original flavor serving as the baseline: dried potatoes, vegetable oil, determinate yellow corn flour, cornstarch, rice flour, maltodextrin, mono- and diglycerides; as are the nutritional profiles. The various flavors have different additives including: lactic acid, yeast extract, cream, citric acid, blue cheese, butter milk, turmeric extract color, onion powder, and Red 40 Lake. It is possible that one or more flavor additives may affect wood-fungus extension rates, and future work could determine any specific impacts of flavor additives on growth rate and reference substrate performance.

From an equipment perspective we believe that slightly-more sophisticated measurement tools may be useful. For example, simple flat-bed document scanners could be used to achieve time-lapse imaging of radial extension to enable more precise quantification of extension dynamics^88^. Additionally, flat-bed scanners have been used to quantify the total number of hyphal tips, total number of hyphal segments, total number of network nodes, total length of the mycelium, growth angle, and branching angle^88^. Furthermore, using time-lapse imaging could speed up determination of extension rates and reduce the size of the plates being used; transitioning from 9 cm plates to 24-well culture plates could enable increased throughput and sample numbers, and better support statistical analysis of differences in strain performance.

Improved protocols that reduce sources of variation which inevitably arise in the making of measurements across many locations are worth exploring. More specifically, the observed variation among labs may be due to differences in the handling of the organisms, the substrate, the cultivation environment, or deviations from the simple experimental protocol supplied. While we advocated for participants to start growth assays immediately upon receipt of our kit, the protocol itself did not prescribe immediate experimentation; although we sent all labs stock cultures from the same mother plate at the same time, some labs were unable to start work immediately and had to store their cultures at 4C until they were able to conduct the experiment, at which time they would have followed the portion of our simple protocol that explained how to prepare starter cultures prior to beginning the experimental plates. One lab requested a second kit after their plates dried out because of prolonged storage; we sent a new kit with samples from our mother plate and provided no further instructions. Fungal colonies age with both time and sub-culturing, and such aging may have an effect on colony radial extension rates. Therefore, prolonged storage at 4C, delays in sampling from stock plates to make starter plates, or delays in sampling from starter plates to make experimental plates may have an effect on radial extension rates and may contribute to the observed variation across labs. It is also possible that the observed variation is due to contamination of the stock, starter, or experimental plates; two labs reported contamination of their kits. One lab was able to clear the contamination and proceed; we observed little variation between their results and other labs. We sent a new kit to the second lab that reported contamination and they proceeded with experiments, reporting the most consistent results. Other potential sources of variation include variation in substrate preparation or timing of colony radius traces. Of note, none of the participating labs had any prior experience working with filamentous fungi. Going forward, we would expect participating labs to become more comfortable working with filamentous fungi and using improved methods for preparing media. Reuse of materials, measures, and methods should itself help with broader adoption of improved metrology practices involving filamentous fungi.

Our community-based measurement methods incentivize practitioners to coordinate the reuse of standard materials, methods, strains, and to share information supporting work with wood-degrading fungi. We hope our approach will realize benefits for reproducible science and reliable fabrication processes. In particular, widely-available reusable materials, measures, and methods that enable practitioners at the edges of biological material supply-chains could help advance mycological production as a distributed manufacturing platform and contribute to a flourishing bioeconomy that benefits all earthlings and the planet.

## MATERIALS and METHODS

### Strains

We acquired four sequenced strains of wood-degrading fungi from various sources (please see SI, Table 1). Upon receipt each culture was expanded to multiple 4C storage plates (9 cm, VWR, Radnor, PA) and culture slants. Culture slants were made by pouring approximately 5-10 mL of PDYA at an angle into a 15 mL Falcon tube (VWR, Radnor, PA) and allowing the PDYA to solidify along the inner sidewall of the tube. Each slant is inoculated with an approximately 2 cm piece of mycelium tissue, para-filmed, and stored in darkness at room temperature.

### Strain Maintenance

All strains were maintained on PDYA by transferring 5mm plugs (Biopunch, Ted Pella, Inc.), from leading edge tissue to fresh PDYA plates for storage at 4C or into slants for storage at room temperature. All starter plates for an organism are inoculated from a common source plate or slant as indicated.

### Pre-culture

Each organism was cultured on PDYA starter plates in darkness at 30C for three days prior to experiments. Starter plates were initiated as indicated by either transferring a 5 mm plug from leading edge tissue of 4C storage plates onto fresh PDYA starter plates or transferring starter tissue from slants onto starter plates.

### Substrates

NIST RM 8492 was obtained from NIST. U.S.-based Pringles™ Original flavor were acquired from Amazon.com, assorted flavors were acquired from Walmart (Bentonville, AR), and Original flavor Pringles™ from the five Pringles™ global production factories were acquired from various retail stores in the countries of origin. Interlaboratory participants acquired locally-sourced Pringles™ from various local retail stores. PDYA was prepared using various laboratory grade reagents (per 0.5 L; 20 g glucose, Sigma; 10 g starch from potato, Sigma; 7.5 g Yeast Extract, EMD Millipore Corporation; 5.0 g Bacto Agar, Becton, Dickinson and Company). Yeast Synthetic Defined (YSD) media was prepared according to the manufacturer recommendations (Takara Bio USA, Inc.). Commercially available woodchip substrate was purchased from Out Grow, IL, USA. Hardwood and Softwood pallets were obtained from All Good Pallets, Inc. (Newark, CA) and chipped to 1-2 cm particle size in a batch process prior to preparation for experiments. Cardboard was harvested from Amazon.com boxes in the Shriram Center for Bioengineering and Chemical Engineering recycle bins.

### Substrate Preparation

Agar-based aqueous extract experimental plates were made for each substrate. Each solid substrate was weighed, ground for 1.5 mins in a Magic Bullet (Walmart, Bentonville, AR), and the ground substrate was suspended in deionized water in a glass beaker to make 2% aqueous extract solutions. Aqueous extractions were performed by boiling the substrate and deionized water mixture in an autoclave for 45 minutes at 121C. The aqueous extract was filtered under vacuum through a funnel plugged with cheese cloth (Labscientific, Inc.) into an Erlenmeyer flask. Solid particulate was discarded and the liquid broth was harvested to make agar-based plates. Agar was added to the aqueous extract broth to make 1% agar solutions for all substrates. The aqueous extract broth and agar mixture was autoclaved at 121 C for 30 minutes prior to use, allowed to cool, and 20 mL of the mixture was dispensed into each plate.

### Plate-based Colony Radial Extension Measurements

Plate-based measurements were conducted in triplicate. Experimental plates were inoculated with a 5 mm plug of leading edge tissue from starter plates and incubated in the dark, at 30C, in ambient humidity (Figure 1: Compost, Manure, Woodchip, PDYA, YSD) or approximately 80% humidity (Figure 1: NIST RM 8492; Figures 2 – 7), until the fastest growing fungus reached the edge of the plate (approximately four days for *Phanerochaete chrysosporium* on PDYA). Every 24 hours the leading edge of each colony was traced with a marker. Measurements of distances between all colony leading edge traces were performed on the final day (Figure 1). Measurements were obtained by hand using a ruler (Figure 1: Compost, Manure, Woodchip, PDYA, YSD) or using image analysis software, as indicated (Figure 1: NIST RM 8492; Figures 2-7). Measurements were recorded in a .CSV file and used for downstream analysis.

### Interlaboratory Study Measurement Kit

Each participating laboratory was provided a measurement kit (SI, Figure 2). Kits contained four fungi of interest, a coring tool for excising fungal tissue plugs, cheese cloth for filtering aqueous extracts, and centrally-provided Pringles substrate. The contents were packaged within the original product packaging and shipped via FedEx to participating labs. Simple instructions for conducting colony radial extension measurements were emailed to participants. Briefly, participants were instructed to culture their starter plates accordingly for three days prior to starting experimental plates, perform colony radius traces for five days, image their plates, and email the images to our group. Interlaboratory study images were processed identically to all other experimental images.

### Image Analysis

Plates from experiments performed by our group were imaged using an iPhone 8. Images from our interlaboratory study were acquired by participants using their preferred method and returned to us for image processing and analysis. Images were converted to .PNG using the Mac application Preview, measurement axis were superimposed (Figure 1), and the images were subsequently analyzed in Fiji (ImageJ)^89^. The Fiji Set Scale tool was used to calibrate pixel distance to millimeters using the Petri dish diameter as a reference distance. The Measurement tool was used to measure distances between traces along predetermined superimposed axis starting from a central point on the edge of the inoculation plug (Figure 1). Values were recorded in a spreadsheet for downstream analysis.

### Statistical Analysis

All analysis was performed using custom Python scripts. The Pandas library was used for data curation^90^. Where appropriate contaminated plate data was dropped from the dataset (Figure 5: Original-USA, two plates dropped due to contamination). Normality of data was checked using the Shapiro-Wilk test in the StatsModels module^91^. StatsModels ANOVA was used to test for significant differences within factors. Tukey HSD in StatsModels was used to identify statistically significant differences between groups. The Scikit-learn library used for feature normalization and linear regression models^92^. All plots were made using MatPlotLib and Seaborn libraries^93^.

## ACKNOWLEDGMENTS

We thank Richard M. Murray for connecting us to his Caltech research group, Samuel Clamons and Andrew Halleran for gathering interlaboratory data, Han Wosten for providing us with a sample of *Schizophyllum commune*, Phil Ross and Kabir Peay for discussions and advice, Jonathan Calles for discussions and manuscript edits, Jensina Froland, Eden Grown-Haeberli, and Isaac Justice for help with pilot experiments, Cimeron Morrissey for providing Pringles™ from Malaysia, Pawel Kusmierski for acquiring Pringles™ from Poland, and Lin Feng for acquiring Pringles™ from China. Patrick Archie for providing compost from the Stanford Farm.

## ATTRIBUTIONS

R. P. and D.E. conceived of the project, wrote, and edited the manuscript. R.P. performed the experiments and image and statistical analysis. M.L. performed pilot and commonly-used biomass experiments. N.G., F.S., S.C., R.K. performed interlaboratory experiments. M.L., N.G., F.S., S.C., R.K., K.A., P.S. edited the manuscript.

## IMAGE ATTRIBUTIONS

Images in table S1 were acquired from Google Images with the “Labeled for reuse” filter. The image of *S. commune* is in the public domain and was taken by Bernard Spragg; of *T. versicolor* under Creative Commons License BY 2.0 by Maja Dumat, of *G. lucidum* by Eric Steinert under Cc-by-sa-3.0,2.5,2.0,1.0, and of *Phanerochaete Spp.* by James Lindsey under Creative Commons license SA 3.0 unported and 2.5 Generic.

## Supplementary Information

**Figure S1.**
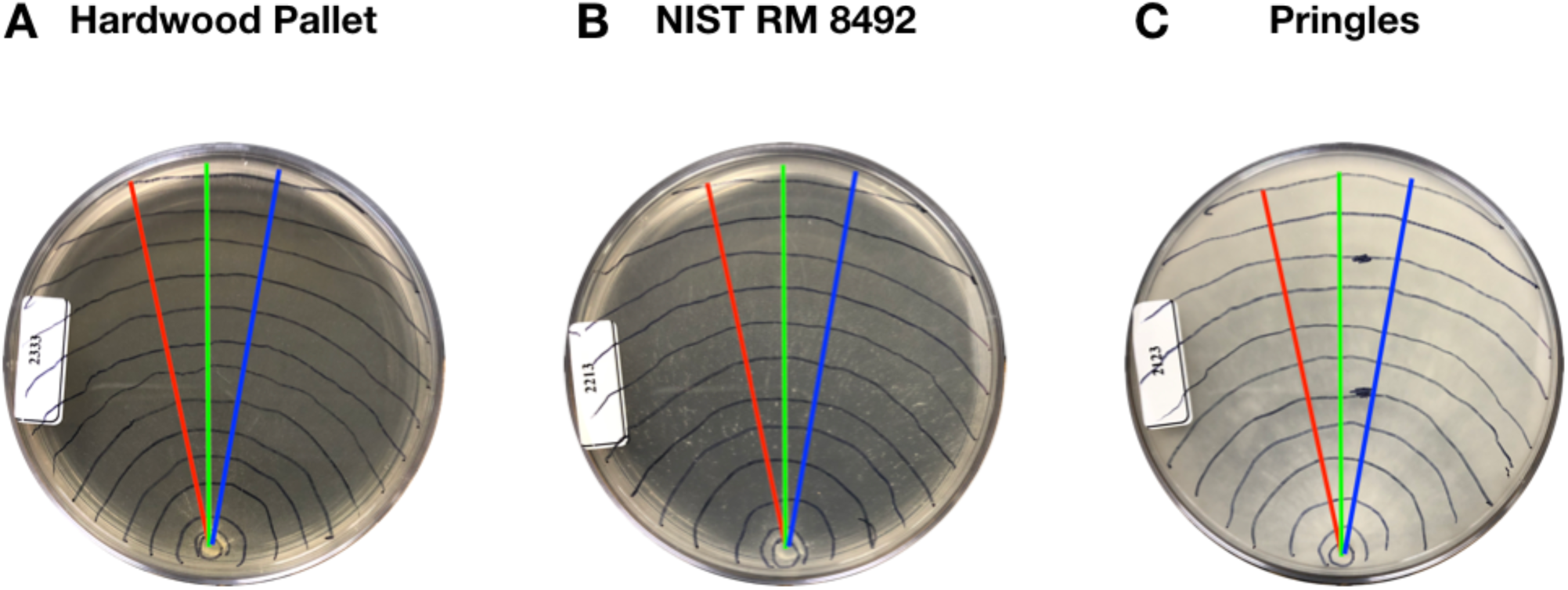
Plate-based assays enable characterization of various parameters of filamentous fungi physiology. In addition to radial extension, density of aerial hyphae, pigmentation, branch rate, and other important characteristics of filamentous fungi can be qualitatively and qualitatively determined using plate-based assays. We observed qualitative differences in mycelium density and aerial hyphae density depending on the substrate composition. Pictured here are representative images of *G. lucidum*.

**Figure S2.**
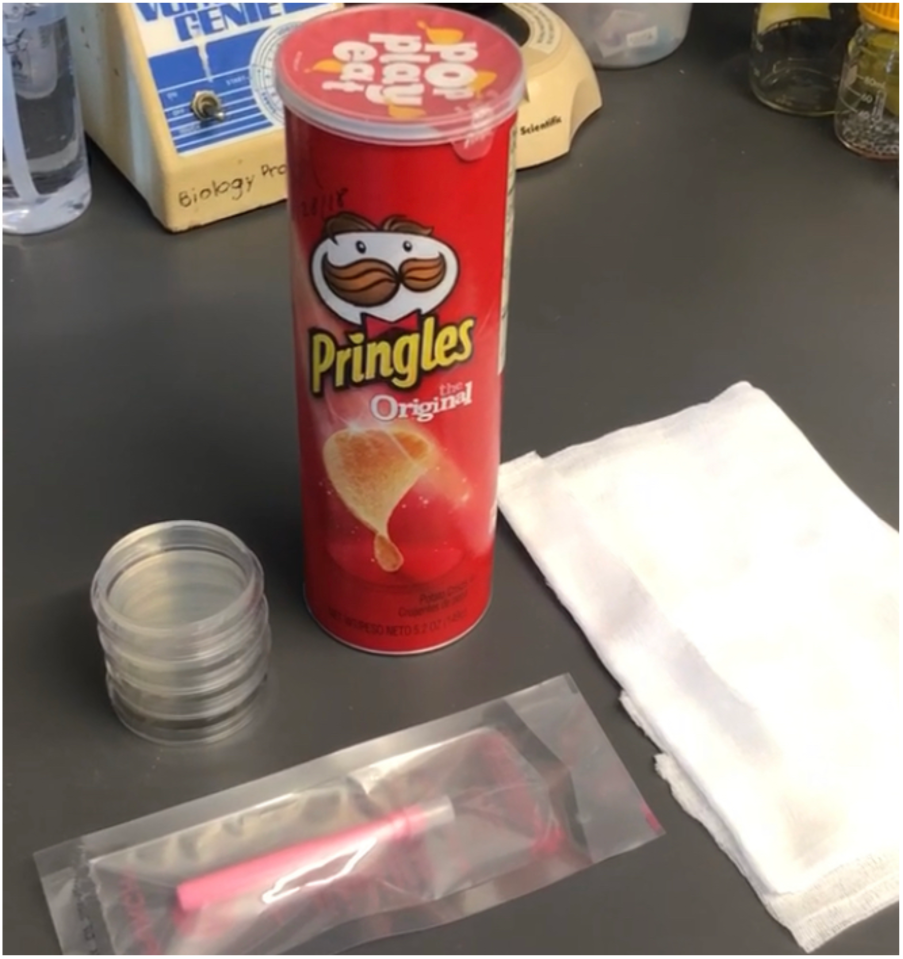
Myco-Metrology Kit v1. We made kits containing centrally-provided Pringles™, fungal cultures, a coring tool, and cheese cloth.

**Table S1.**
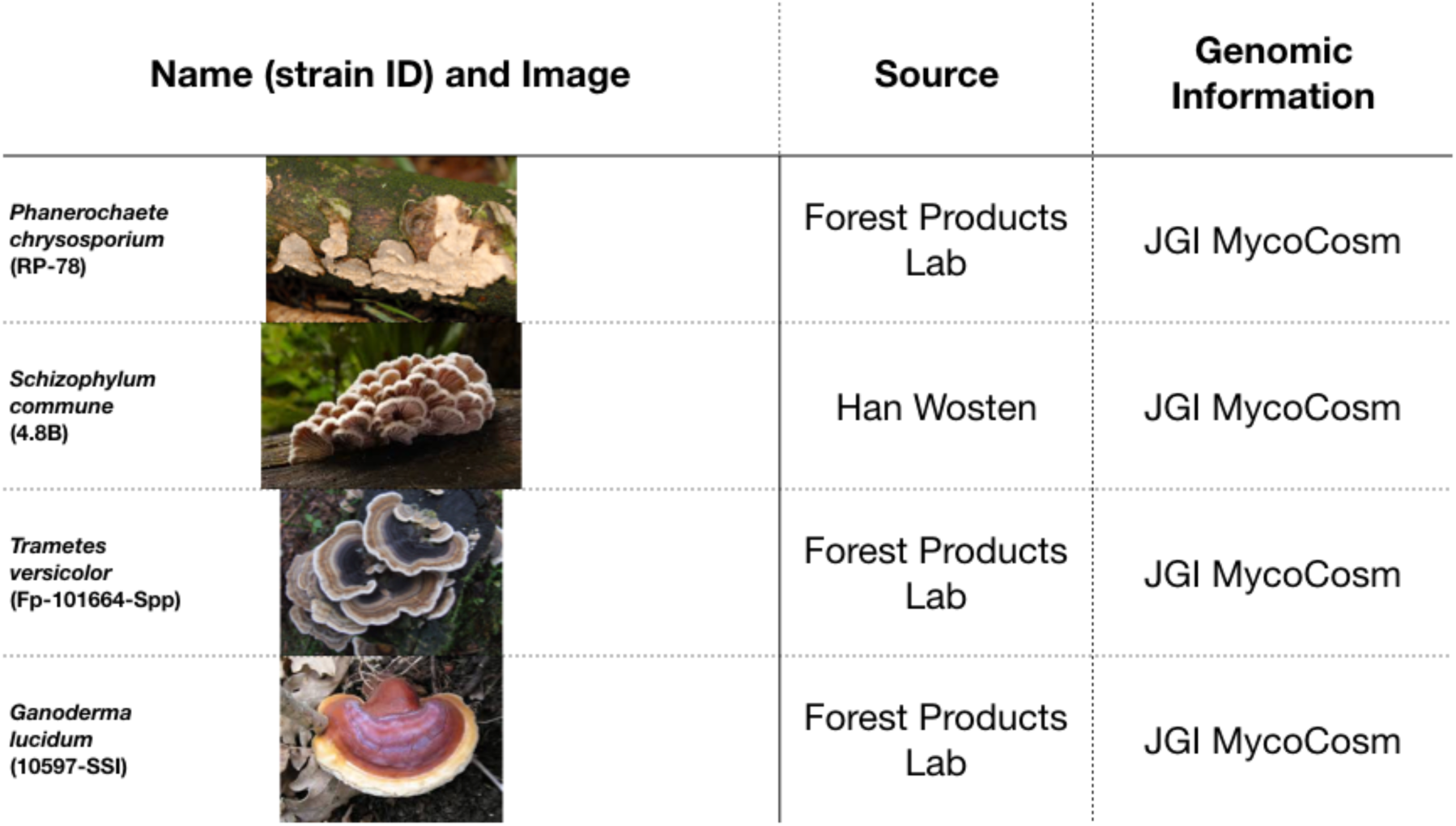
**Organisms used in this study.** As noted.

| Name (strain ID) and Image |  | Source | Genomic Information |
| --- | --- | --- | --- |
| *Phanerochaete chrysosporium* (RP-78) |  | Forest Products Lab | JGI MycoCosm |
| *Schizophyllum commune* (4.8B) |  | Han Wosten | JGI MycoCosm |
| *Trametes versicolor* (Fp-101664-Spp) |  | Forest Products Lab | JGI MycoCosm |
| *Ganoderma lucidum* (10597-SSI) |  | Forest Products Lab | JGI MycoCosm |

**Table S2.**
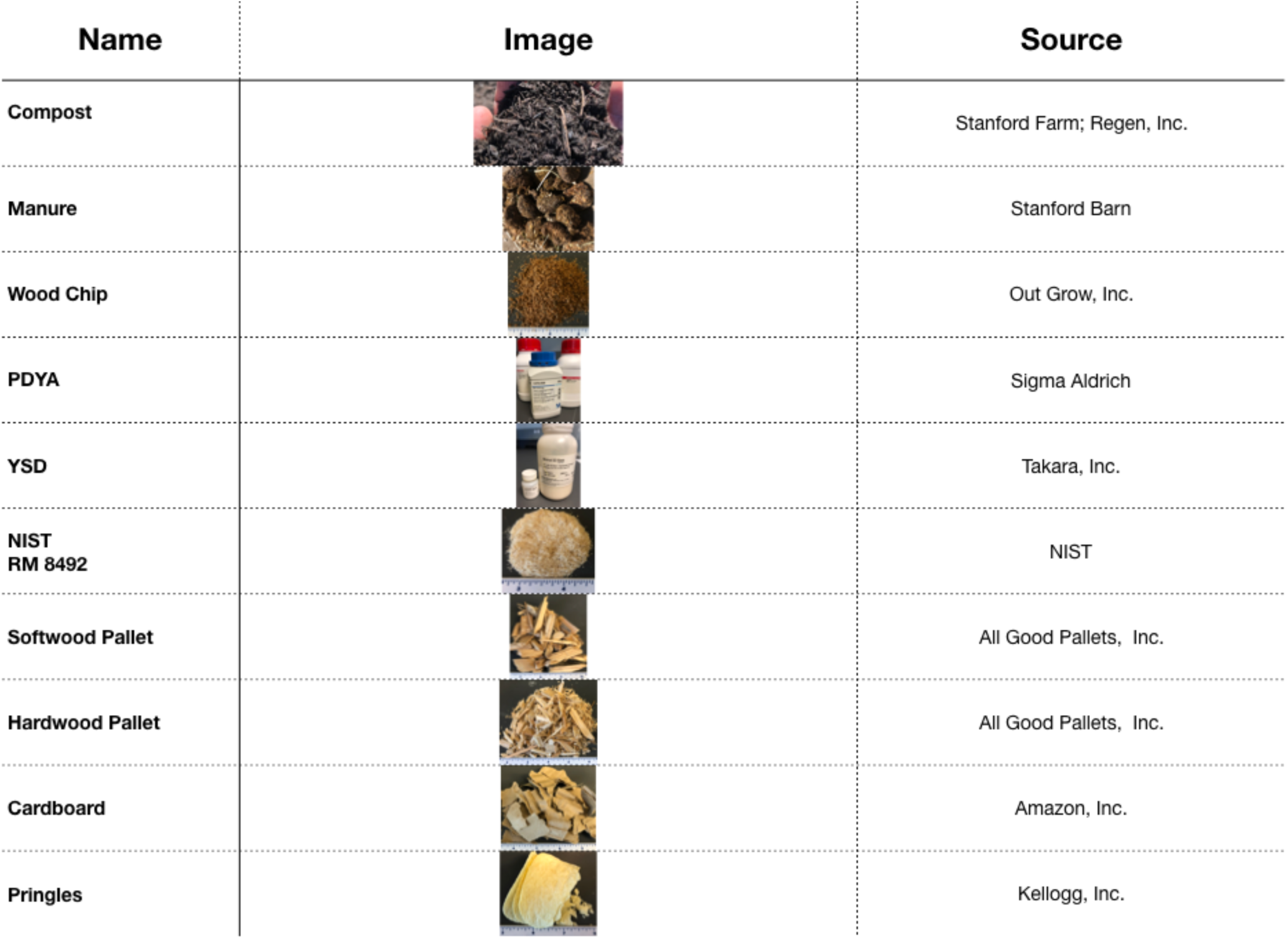
**Substrates used in this study.** We sourced various substrates that are either commonly-used, laboratory-grade, fully-traceable, or widely-available.

| Name | Image | Source |
| --- | --- | --- |
| Compost |  | Stanford Farm; Regen, Inc. |
| Manure |  | Stanford Barn |
| Wood Chip |  | Out Grow, Inc. |
| PDYA |  | Sigma Aldrich |
| YSD |  | Takara, Inc. |
| NIST<br>RM 8492 |  | NIST |
| Softwood Pallet |  | All Good Pallets, Inc. |
| Hardwood Pallet |  | All Good Pallets, Inc. |
| Cardboard |  | Amazon, Inc. |
| Pringles |  | Kellogg, Inc. |

**Table S3.** **Reference Mass Fraction Values for Constituents in RM 8492.** The entire table and all values shown here are reproduced directly from the NIST Report of Investigation for Reference Material 8492^84^. NIST RM 8492 Eastern Cottonwood Whole Biomass Feedstock is fully-traceable and compositionally-characterized candidate reference material. RM 8492 is intended primarily for use in evaluating analytical methods for the determination of summative composition of lignocellulosic material. The RM can also be used for quality assurance when assigning values to in-house control materials.

|  | Mass Fraction<br>(%) | Expanded Uncertainty<br>(%) | Coverage Factor, *k* |
| --- | --- | --- | --- |
| Water Extractives | 2.9 | 1.2 | 2.20 |
| 95 % Ethanol Extractives (after water extraction) | 1.54 | 0.63 | 2.20 |
| Sucrose<sup>(b)</sup> | 0.045 | 0.046 | 2.20 |
| Whole Ash | 0.96 | 0.23 | 2.18 |
| Extractives-Free Ash | 0.741 | 0.077 | 2.26 |
| Glucan | 44.6 | 1.2 | 2.18 |
| Xylan | 13.39 | 0.39 | 2.18 |
| Arabinan | 0.35 | 0.30 | 2.18 |
| Galactan | 0.55 | 0.53 | 2.20 |
| Mannan | 2.16 | 0.30 | 2.18 |
| Structural Sugars | 61.0 | 1.8 | 2.18 |
| Total Lignin<sup>(c)</sup> | 27.2 | 1.3 | 2.18 |
| Acid-Insoluble Residue | 24.0 | 1.2 | 2.20 |
| Acid-Soluble Lignin | 2.2 | 1.7 | 2.20 |
| Acetyl | 3.3 | 1.8 | 2.23 |
| Nitrogen | 0.17 | 0.11 | 2.31 |
| Total Component Closure<sup>(d)</sup> | 99.4 | 1.5 | 2.26 |

